# SARS-CoV-2 infection induced alterations in ADAR editing patterns differ between patients who developed critical compared to non-critical COVID-19

**DOI:** 10.1101/2025.10.22.684004

**Authors:** Aiswarya Mukundan Nair, Helen Piontkivska

**Affiliations:** Department of Biological Science, Kent State University, Kent, OH, USA; Brain Health Research Institute, Kent State University, Kent, OH, USA

**Keywords:** SARS-CoV-2, Severe acute respiratory syndrome coronavirus 2, RNA editing, ADAR editing, transcriptome, critical COVID-19

## Abstract

COVID-19, caused by the SARS-CoV-2 virus, has a wide spectrum of clinical presentations even among individuals with similar demographics. Disease severity has been linked with viral recognition-triggered expression of Interferons (IFNs) and Interferon stimulated genes (ISGs). Among these ISGs, ADARp150 is a member of adenosine deaminases acting on RNA (ADARs) enzyme family. ADARs are RNA editing enzymes that contribute to transcriptome diversity and modulate immune response during viral infections. While previous studies have identified altered ADAR expression and editing patterns during SARS-CoV-2 infection, it remains unknown whether ADAR expression and activity differ between patients with varying severities of COVID-19, specifically, in individuals who developed critical compared to non-critical COVID-19. We address this question by analyzing a publicly available, deeply sequenced whole blood RNA-seq dataset from individuals with either critical or non-critical COVID-19, matched for age, sex, and presence of comorbidities. Our results show differential expression of thousands of genes, including those involved in neutrophil degranulation, and upregulation of ADAR1 and its isoform ADARp110 in patients with critical COVID-19. We further identify global differences in the total number of edits, driven by ADAR1 and ADAR2 expression levels in critical but not in non-critical patients. ADAR activity also differed within Alu elements and in the proportion of edits with varying functional consequences. We further identified severity specific editing events, including nonsynonymous edits, within distinct biological pathways. Moreover, we identified 140 high confidence editing sites within 126 genes, that are differentially edited between the two patient groups. These genes were enriched in infectious disease, cell cycle, signal transduction, RNA and protein metabolism in addition to inflammatory pathways such as neutrophil degranulation and signaling by interleukins. Enrichment/modulation of neutrophil degranulation pathway at transcriptional and post transcriptional levels suggest the importance, complex regulation, and contribution of this pathway in COVID-19 disease severity. Finally, using a random forest classifier, we identified a set of differentially edited sites that could serve as molecular markers for COVID-19 disease severity. Together, our study demonstrates varying expression and editing patterns of ADARs between critical and non-critical patients, suggesting a potential role of ADAR editing in varying severity of COVID-19 pathogenesis.

## Introduction

Coronavirus disease 2019 (COVID-19), caused by the betacoronavirus SARS-CoV-2, is characterized by a range of clinical severities even among individuals with similar demographics (C. Huang et al., 2020; Lowery et al., 2021). Accordingly, SARS-CoV-2 infections are categorized into asymptomatic, mild, moderate, severe, and critical illnesses (CDC, 2024; Montenegro et al., 2024). A majority of SARS-CoV-2 infections result in asymptomatic or non-critical COVID-19 with influenzae-like symptoms such as headache, fever, cough, myalgia, fatigue, shortness of breath, and sore throat; however, a smaller proportion may progress to critical, life-threatening diseases (G.-U. Kim et al., 2020; Sauerwald et al., 2022). Patients with critical COVID-19 often present with pneumonia, acute respiratory distress syndrome (ARDS), and/or multi-organ failure requiring external respiratory support, intensive care, which are often associated with high mortality rates (Gebremeskel et al., 2024; Goh et al., 2020; Guan et al., 2020; Oran & Topol, 2020; D. Wang et al., 2020). Disease severity is associated with a range of host factors, such as age (Davies et al., 2020; Y. Liu et al., 2020), sex (N. Chen et al., 2020; Meng et al., 2020; Takahashi et al., 2020), presence of comorbidities (Biswas et al., 2020), blood group status (Ellinghaus, 2023; N. Liu et al., 2021), presence of neutralizing autoantibodies (Bastard et al., 2020; Chauvineau-Grenier et al., 2022; Koning et al., 2021; Troya et al., 2021), and genetic variants (Niemi et al., 2021; Pathak et al., 2022; Severe Covid-19 GWAS Group et al., 2020). Additionally, during active infection, a highly dysregulated innate immune response - consisting of impaired production of Interferons (IFNs) and interferon stimulated genes (ISGs), along with excessive NF-kB driven inflammatory response - is a key feature distinguishing patients with critical compared to non-critical COVID-19 (Del Valle et al., 2020; Hadjadj et al., 2020; Merad et al., 2022; Smith et al., 2022).

SARS-CoV-2 is a single stranded, positive sense RNA virus (V’kovski et al., 2021). Upon infection, viral pathogen-associated molecular patterns (PAMPs), including replication intermediaries, are detected by the innate immune receptors (Y.-M. Kim & Shin, 2021; S. Zhang et al., 2022). Viral recognition triggers a downstream antiviral cascade resulting in the production of IFNs and ISGs. There are three main types of IFNs, namely, type I (IFN-α/β), type II (IFN-γ), and type III (IFN-λ) (Lazear et al., 2019). These IFNs bind to interferon stimulated response elements (ISREs) to induce transcription of interferon stimulated genes (ISGs), that act through direct and indirect mechanisms to establish an antiviral state (Ivashkiv & Donlin, 2014; Minkoff & tenOever, 2023). Genetic variants in type I and III IFN pathway genes and autoantibodies against type I IFNs (IFN-α2 and IFN-ω) have been observed in patients with critical COVID-19 (Bastard et al., 2020; Koning et al., 2021; Qian Zhang et al., 2020). Furthermore, IFN levels vary according to disease severity and delayed or impaired production of type I (characterized by low levels and activity of IFN-α and absence of IFN-β), III IFNs and ISGs, is observed in patients with critical disease (Blanco-Melo et al., 2020; Galani et al., 2021; Hadjadj et al., 2020). However, contrasting studies also exist where critical COVID-19 patients displayed increased levels of IFNs and ISG expression (Broggi et al., 2020; Lucas et al., 2020). These differences observed in the levels of IFNs and ISGs from different studies could be attributed to the multiple factors, including differences in population studied, anatomical site being studied, differences in the subtype of IFN-α reported, and differences in assays used to measure the levels of cytokines (technical approaches) (Smith et al., 2022). Nevertheless, differences in expression of IFNs and ISGs is widely observed between patients who develop critical compared to non-critical COVID-19, underscoring the importance of IFNs and their downstream effectors in determining COVID-19 severity (Smith et al., 2022).

Among hundreds of ISGs produced in response to SARS-CoV-2 infection is Adenosine Deaminase Acting on RNA 1 (ADAR1), specifically, the longer cytoplasmic isoform ADARp150, whose promoter region incorporates an ISRE. This ADAR edits RNA molecules post transcriptionally to convert adenosines (A) to inosines (I), within double stranded RNA (dsRNA) regions. These substitutions are then interpreted as A-to-G substitutions by cellular machinery, including elements involved in translation. Thus, ADAR mediated editing is a key player in the dynamic regulation of gene expression and proteomic diversity through introduction of recoding sites within proteins or through its effect on various regulatory mechanisms such as alternative splicing, microRNA biogenesis and targeting (Hsiao et al., 2018; Tomaselli et al., 2013). In addition to ADARp150, ADAR1 also exists as a constitutively expressed, predominantly nuclear isoform, ADARp110 (Savva et al., 2012a). It is noteworthy that ADARp110 is coexpressed with ADARp150 due to leaky ribosomal scanning downstream of the ADARp150 start codon (Sun et al., 2021a). Additionally, increased levels of ADARp110 have also been observed during SARS-CoV-2 and other viral infections (Nair & Piontkivska, 2025; Tariq & Piontkivska, 2024). In addition to ADAR1 (ADAR), the human genomes encode for two other ADAR genes, ADAR2 (ADARB1) and ADAR3 (ADARB2). While most viral infections are associated with changes in the expression and activity of ADAR1 and its isoforms, alterations in ADAR1 expression can also affect the expression and activity of other ADARs, such as through competition for substrates (Savva et al., 2012a) and formation of heterodimers (Cenci et al., 2008).

ADARs edit both viral and host transcripts, and this editing could shape the outcomes for both the virus and the host (Di Giorgio et al., 2020; Piontkivska et al., 2021; Tariq & Piontkivska, 2024). Mechanistically, it was demonstrated that the SARS-CoV-2 nucleocapsid (N) protein directed viral RNAs to ADARs (ADAR1/2), within infection induced stress granules to promote viral editing (Li et al., 2024). Accordingly, widespread ADAR editing signatures are observed in the SARS-CoV-2 genome (Di Giorgio et al., 2020; Picardi et al., 2022). Deep transcriptomic studies have identified a higher proportion of ADAR edits in the minor viral RNA population compared to the consensus viral population suggesting an inverse correlation between ADAR editing and viral load (Ringlander et al., 2022). Multiple nonsynonymous ADAR edited sites have also been detected in the receptor binding motif of SARS-CoV-2 spike protein, causing structural changes that alter viral binding to host receptors, potentially affecting viral infectivity (Ringlander et al., 2022; Song et al., 2022). Host ADAR mediated editing within the SARS-CoV-2 genome has been proposed as a major contributor to viral mutation, infectivity, transmissibility, fitness, and evolution during the pandemic (Azgari et al., 2021a, 2021b; Di Giorgio et al., 2020; Kosuge et al., 2020; Mourier et al., 2021; Picardi et al., 2022; R. Wang et al., 2020).

Within the host, ADAR edits are found in both coding and non-coding regions (Gabay et al., 2022; Nishikura, 2016). Editing in these regions plays a crucial role in mediating immune responses, preventing autoimmunity, inflammation, and regulating gene expression (Piontkivska et al., 2021; Q. Wang et al., 2017; Yuan et al., 2023). As mentioned above, editing within protein coding regions can result in non-synonymous substitutions that may alter the structure and function of the encoded protein, thereby increasing proteomic diversity from a limited set of genes (Eisenberg & Levanon, 2018; Hood & Emeson, 2012; Rosenthal & Seeburg, 2012). Many such highly conserved sites with significant physiological and pathological impacts have been identified in genes associated with cancer, neuronal function, and cardiovascular health (Hu et al., 2015; Jain et al., 2018; Pinto et al., 2014). However, the majority of ADAR editing occurs in the non-coding regions of transcripts, such as primate specific Alu retrotransposons, that can form dsRNAs that are primarily located within the 3’UTR and intronic regions (D. D. Y. Kim et al., 2004; Solomon et al., 2017). Editing reduces their dsRNA characteristics and recognition by endogenous immune sensors, thereby preventing autoimmunity (Jiao et al., 2024; Liddicoat et al., 2015). Notably, proinflammatory responses due to altered ADAR editing is a characteristic of several inflammatory diseases, including atopic dermatitis (AD) (Karmon et al., 2023), multiple sclerosis (MS) (Tossberg et al., 2020), Aicardi Goutières Syndrome (AGS) (Chung et al., 2018), and human inflammatory bowel disease (Aune et al., 2022). A similar phenotype of reduced levels of ADAR editing within endogenous Alu elements has been observed in SARS-CoV-2 infected human lung cells and in lung biopsies of COVID-19 patients. In these cases, accumulation of unedited Alu RNAs was associated with activation of IRF and NF-kB driven transcriptional responses, potentially contributing to COVID-19 associated inflammation (Crooke et al., 2021b, 2021a). Similarly, changes in ADAR editing were also observed in different ocular tissues during SARS-CoV-2 infection and were linked to ocular manifestations of COVID-19 (Jin, Liang, Huang, et al., 2024). We have previously identified increased global ADAR editing levels mid SARS-CoV-2 infection that were found to persists in a subset of individuals, even after viral clearance, in patients with mild COVID-19 (Nair & Piontkivska, 2025). In a recent study, Chattopadhyay et al. (2024) identified differences in ADAR editing within host long noncoding RNAs (lncRNAs) among patients infected with Delta, PreVoC, or Omicron variants, with most distinct editing patterns observed during infections with Delta variant. Their findings suggest that lncRNA editing could be variant specific and may contribute to differences in disease severity observed between these variants (Chattopadhyay et al., 2024). Additionally, differences in host ADAR editing have also been observed in response to COVID-19 vaccinations, where these changes have been proposed to have a protective role by modulating host immunity against SARS-CoV-2 infections (Jin, Liang, Pan, et al., 2024). Findings from these studies, along with previous studies from other viral infections (Piontkivska et al., 2019, 2021; Tsivion-Visbord et al., 2020), establish RNA editing as a mechanistic link between viral infections and symptoms observed both during and post viral infections.

While it is well established that host ADAR expression and editing is altered during SARS-CoV-2 infections (Crooke et al., 2021a; M. Huang et al., 2024; Merdler-Rabinowicz et al., 2023; Nair & Piontkivska, 2025), it remains unknown whether the expression and activity of ADARs differ in individuals who develop critical compared to non-critical COVID-19. To address this gap, we analyzed deeply sequenced whole blood RNA sequencing data from a young patient cohort, excluded for major comorbidities, from Carapito et al. (PRJNA722046) (Carapito et al., 2022) to identify differences in ADAR expression and activity, measured as differences in ADAR editing, between patients who developed critical compared to non-critical COVID-19.

## Results

### Expression of RNA editing enzyme ADAR1 is higher in patients with critical COVID-19

To investigate differences in gene expression, including whether the expression of ADAR genes and isoforms differ between patients who developed critical compared to non-critical COVID-19, differential gene expression analysis was performed on whole blood RNA sequencing data using the DESeq2 package (Love et al., 2014). Principle component analysis (PCA) on VST (Variance Stabilizing Transformation) normalized raw counts data showed that PC 1 accounted for 19% variance and separated non-critical from critical patients (Supplementary Figure 1A). After filtering out genes that were expressed at low levels (keeping those with ≥10 reads in ≥90% of samples), 4111 genes were identified as differentially expressed between the two patient groups, using the filtering criteria of log_2_Fold Change ≥ |0.58| (Fold Change ≥ 1.5) and an adjusted p-value < 0.05 (Fig. 1A; Supplementary Table 1A). Within the differentially expressed genes, 2802 genes were upregulated (Supplementary Table 1B), while the 1309 genes were downregulated in critical compared to non-critical patients (Supplementary Table 1C).

**Figure 1:**
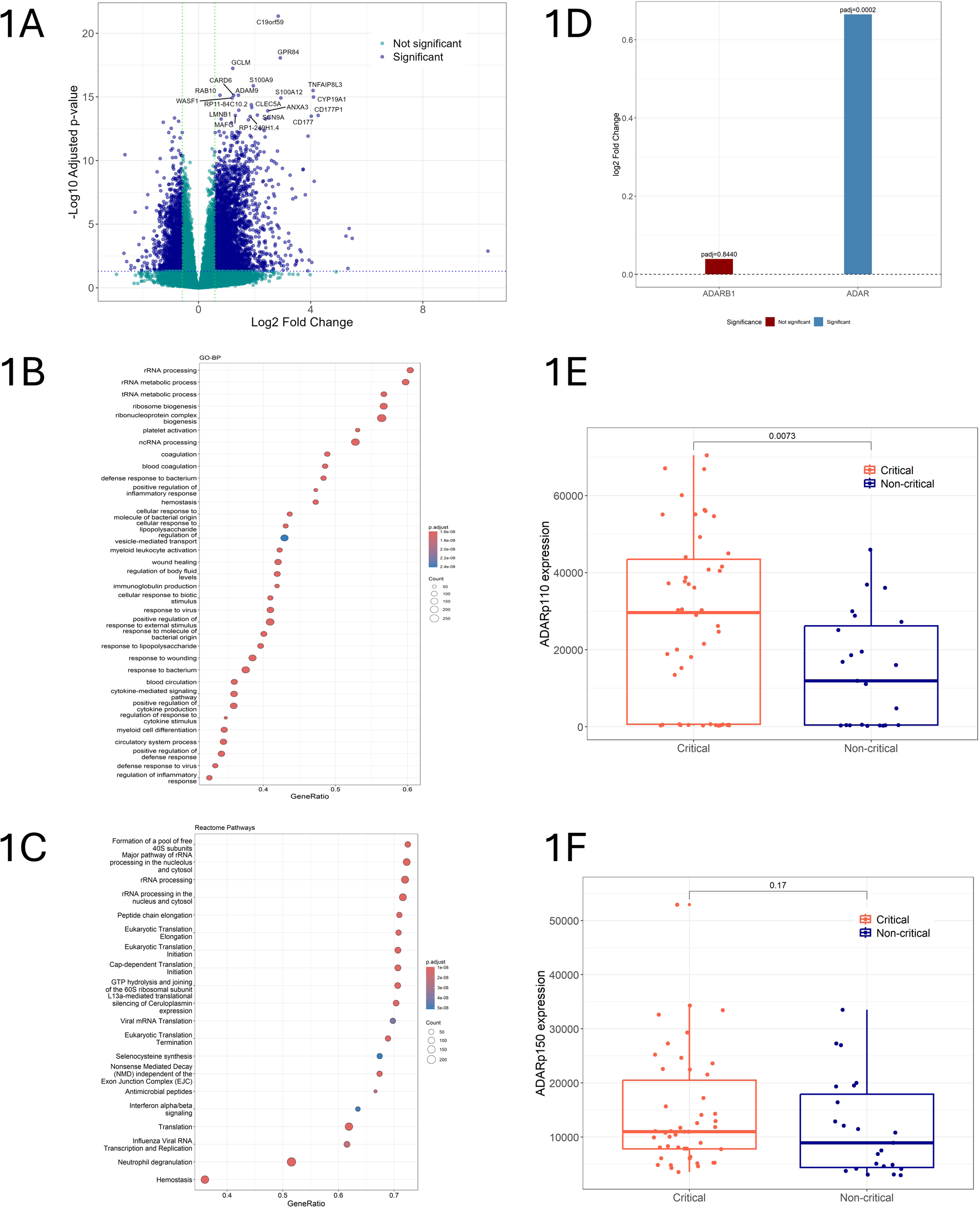
Differences in gene expression, including changes in the expression of ADARs and ADAR1 isoforms in patients with critical compared to non-critical COVID-19. (A) Volcano plot showing differentially expressed genes (log_2_Fold Change ≥ |0.58| and an adjusted p-value < 0.05), with a higher number of genes upregulated in patients with critical COVID-19. Blue dots represent genes with significant difference in expression between the two patient groups. The top 20 differentially expressed genes are labeled on the plot. GSEA analysis of differentially expressed ranked by Log2FC with gene sets from (B) GO – Biological Processes (BP) (C) Reactome pathway. (D) DESeq2 analysis results for ADAR genes: ADAR1 (ADAR) and ADAR2 (ADARb1). ADAR1 is significantly higher in critical (Log2FC = 0.6647and padj = 0.0002) compared to non-critical patients. While expression of ADAR2 (Log2FC = 0.03939 and padj = 0.8440) is minimally different between critical and non-critical patients, this difference was not significant. DESeq2 normalized expression of ADAR1 isoforms: ADARp110 (E) and ADARp150 (F). ADARp110 expression is significantly upregulated in critical patients (Wilcoxon rank-sum test, p-value 0.0073) whereasADARp150 expression was minimally higher in critical patients compared to non-critical patients (Wilcoxon rank-sum test, p-value 0.17).

To understand which biological processes and pathways experienced differential gene expression, we performed Gene Set enrichment Analysis (GSEA) using gene list ranked by Log2FoldChange, with gene sets from GO-BP and Reactome. Genes upregulated within critical COVID-19 patients showed enrichment for innate immune system pathways/processes including “neutrophil degranulation”, “interferon alpha/beta signaling”, “antimicrobial peptides”, “cellular response to type I interferon”, hemostasis pathways including “response to elevated platelet cytosolic Ca2+”, “platelet degranulation”, neuronal pathways including “GABA receptor activation”, “inwardly rectifying K+ channels”, and disease pathways including “disease associated with the TLR signaling cascade”. Genes downregulated within critically ill patients were enriched in RNA and protein metabolism pathways/processes such as “formation of a pool of free 40S subunits”, “MHC class II protein complex assembly” and “Nonsense Mediated Decay (NMD) independent of the Exon Junction Complex (EJC)”. The top 20 significant GO-BP and Reactome pathways (padj < 0.05) are shown in the figure 1B and 1C, full list of enriched pathways/GO processes, ordered by the Nominal Enrichment Score (NES) and padj < 0.05 are provided in the Supplementary Table 1D and 1E.

While previous studies have identified elevated expression of ADAR1 in response to SARS-CoV-2 infections (Nair & Piontkivska, 2025; Song et al., 2022), it is unknown whether this increase in expression is consistent or differs between patients with varying disease severity. We found significant differences in ADAR1 expression (Log2FC = 0.6647and padj = 0.0002) between the two patient groups with higher expression observed in patients with critical COVID-19. Furthermore, ADAR2 expression, although without significance, was found to be nominally higher (Log2FC = 0.03939 and padj = 0.8440) in critical patients. While ADAR3 expression was not detected in our samples, this could be due to restricted expression of ADAR3 outside brain tissues (Melcher et al., 1996; Raghava Kurup et al., 2022; Ashley et al., 2024) (Fig. 1D; Supplementary Table 1F). Transcript level expression of constitutively expressed ADAR1 isoform, ADARp110 and IFN inducible isoform, ADARp150 (Rehwinkel & Mehdipour, 2025; Samuel, 2011) were examined as DESeq2 normalized counts. ADARp110 was found to be over expressed in patients with critical illness (Wilcoxon rank-sum test: p value = 0.0073, DESeq2: Log2FC = 0.9388 and padj = 0.2201) (Fig. 1E), while expression levels of ADARp150, though minimally lower in patients with non-critical disease, did not achieve statistical significance (Wilcoxon rank-sum test: p value = 0.17, DESeq2: Log2FC = 0.2905 and padj = 0.5304) (Fig. 1F; Supplementary Tables 1G and 1H).

### ADAR editing patterns differs between patients with critical and non-critical COVID-19

Next, we investigated whether ADAR activity differs between the two patient groups. To access this, we quantified ADAR mediated editing within critical and non-critical patients using two different approaches 1) by comparing the total number of ADAR edited sites and 2) by calculating Alu Editing Index (AEI), as a measure of global levels of ADAR editing within Alu repetitive elements. In agreement with ADAR mediated editing being the most abundant form of substitutions in humans (Eisenberg & Levanon, 2018; Mannion et al., 2015; Walkley & Li, 2017), A-to-G and T-to-C edits (A nucleotides on the complementary strand) representing potential ADAR edited sites were the most abundant form of substitutions within both groups of patients (Fig. 2A, Supplementary Table 2A). However, the average number of both A-to-G and T-to-C substitutions differed between critical and non-critical COVID-19 patients. Specifically, the mean number of ADAR edits per sample were higher in patients with non-critical illness, with an average of 158,022 ± 10,234 edits, while patients with critical illness had an average of 139,285 ± 9307 edits per sample (Supplementary Figure 2 and Table 3). These sites were distributed mainly in intronic (74%), intergenic (19%), 3’UTR (4%), and downstream regions (2%), while a considerable number of sites were also distributed within the exonic regions (1%) (Fig. 2B) and the mean number of editing sites within these regions differed between the two severities (Supplementary Table 2B). Notably, within exonic regions, differences were seen in the number of non-synonymous and synonymous edits between the two patient groups (Fig. 2C). Further, the Alu editing index for A-to-G substitutions was highest among all other substitutions (Fig. 2D). However, unlike the total number of ADAR edits, AEI for ADAR edits within Alu elements was higher in patients with critical COVID-19 (Fig. 2D and Table 3). These differences suggest that ADAR activity may differ between patients who develop critical compared to non-critical COVID-19 in response to SARS-CoV-2 infections.

**Figure 2.**
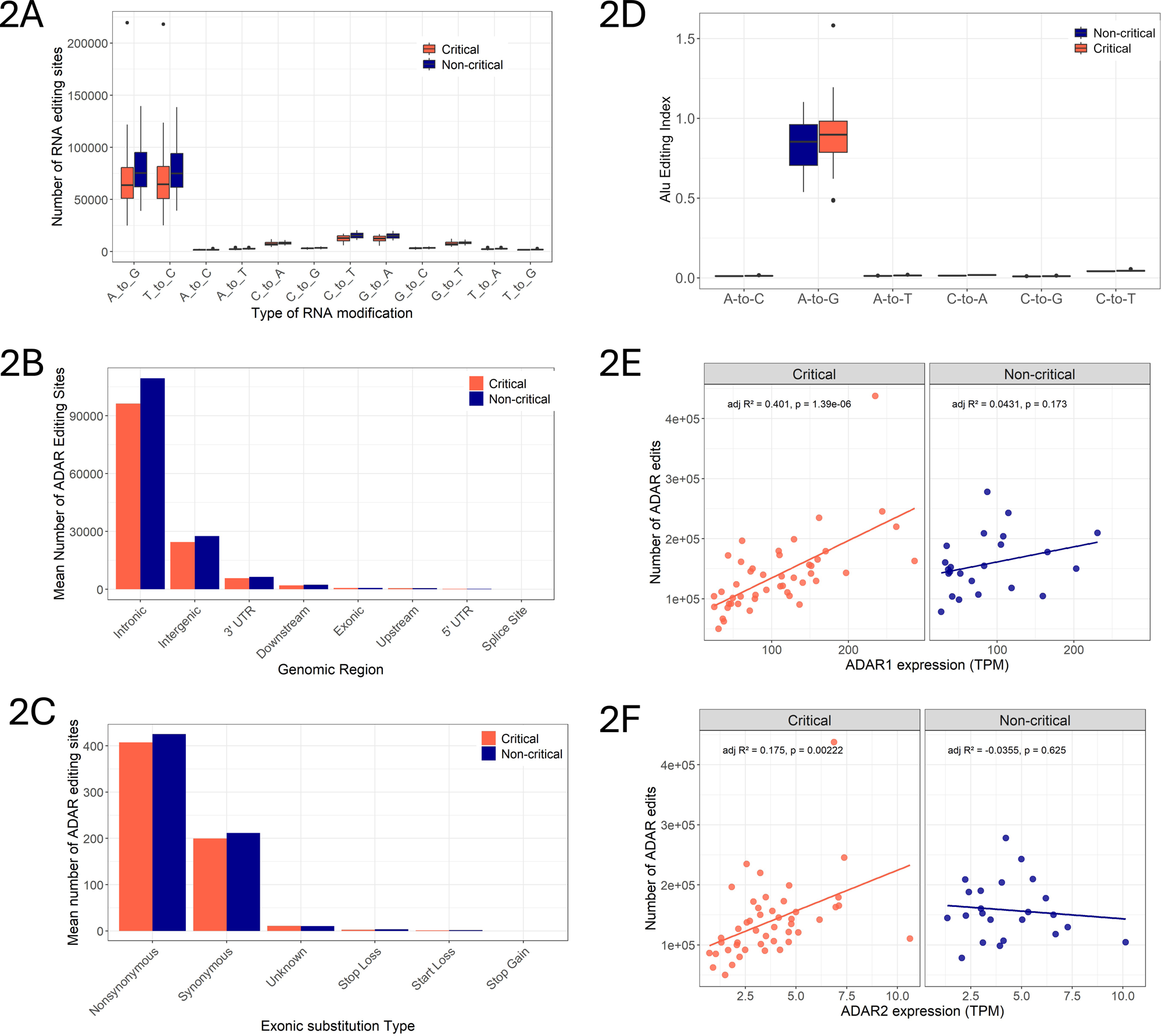
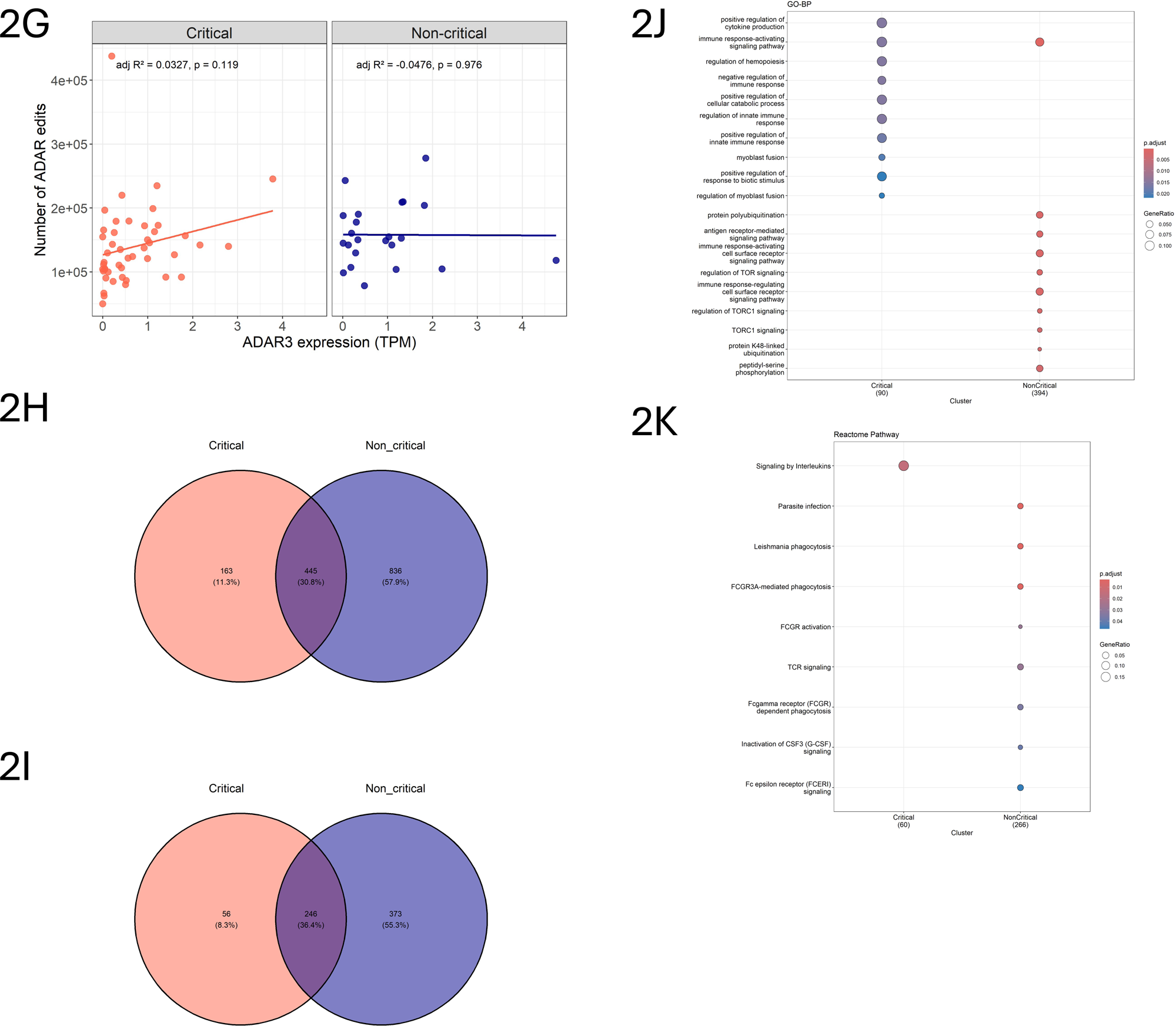
Differences in ADAR editing between critical and non-critical patients. (A) Bar plot shows the number of different substitutions identified by GATK. A-to-G and T-to-C substitutions, representing potential ADAR editing sites, are the most abundant type of substitutions in both groups of patients, however the number of these substitutions are higher among patients with non-critical COVID-19. (B) Distribution of editing sites across genic regions. The average number of edits across all regions was higher for non-critical patients. (C) Number of different exonic substitution types. Non-synonymous substitutions were the most abundant in both groups with higher counts in patients with non-critical COVID-19. (D) Comparison of Alu editing Index (AEI) between patient groups. A-to-G index was higher in patients with critical disease. Correlation between the expression of ADAR1 (E), ADAR2 (F), and ADAR3 (G) measured in TPM, and total number of ADAR edits in critical and non-critical patients. ADAR1 (adj R^2^ = 0.401, p = 1.39 x 10^-6^) and ADAR2 (adj R^2^ = 0.175, p = 0.00222) expressions were significantly positively correlated with the total number of ADAR edits within critically ill patients. Venn diagram comparing the number of ADAR editing sites (H) and corresponding genes (I) that are shared and unique between the two patient groups. Overrepresentation analysis of GO-biological processes (J) and pathways enriched (K) among genes with unique edits in each group. Genes incorporating unique edits were enriched for distinct biological functions.

We further investigated whether the observed differences in total number of ADAR edits between the two patient groups were associated with differences in ADAR expression. Towards this, ADAR expression levels, measured as TPM (Transcripts Per Million) were correlated with the total number of ADAR edits within each patient group. In critical patients, the total number of ADAR edits showed a significant positive correlation with the expression levels of ADAR1 (adj R^2^ = 0.401, p = 1.39 x 10^-6^) and ADAR2 (adj R^2^ = 0.175, p = 0.00222). In contrast, no significant correlation was observed between expression of ADAR1 (adj. R² = 0.0431, p = 0.173) or ADAR2 (adj. R² = –0.0355, p = 0.625) and total editing events in non-critical patients (Fig. 2E, 2F). Similarly, no significant correlations were observed between ADAR3 expression and total editing events in either critical (adj R² = 0.0327, p = 0.119) or non-critical patients (adj R² = –0.0476, p = 0.976) (Fig. 2G), potentially due to negligible expression of ADAR3 in whole blood (Giacopuzzi et al., 2018). We also found ADAR editing sites that were uniquely edited within a severity group, indicating disease severity-specific editing signatures. Specifically, we observed 163 uniquely edited sites within 56 genes in critical COVID-19 patients, while 836 uniquely edited sites within 373 genes in non-critical patients (Fig. 2H, I and Supplementary Table 2D, E). While these uniquely edited sites showed comparable genomic distribution between both groups of patients, with most sites located within the intronic, 3’UTR, and intergenic regions, the number of potential non-synonymous exonic edits varied (Supplementary Figure 2B, C). Of the four exonic edits in NBPF8, SMN1, HSPA1L, and HLA-DRB5 genes, within the non-critical patient group, HLA-DRB5 incorporated a non-synonymous edit at position Chr6: 32487309 and critical patients harbored a non-synonymous substitution within DHRSX gene (position ChrX:2139200). We analyzed the functional consequences of these non-synonymous edits on protein stability using AlphaFold2 (Jumper et al., 2021) structure of HLA-DRB5 and DHRSX in the DDMut tool (Zhou et al., 2023),https://biosig.lab.uq.edu.au/ddmut/, accessed June 06 2025). In critical patients, substitution of Histidine (H) to Arginine (R) at position 292 within DHRX was predicted to result in a ΔΔG of -1.27 kcals/mol (Table 1, Supplementary Figure 2E), while in non-critical patients, the substitution of Serine (S) to Glycine (G) at position 164 within HLA-DRB5 was predicted to result in a ΔΔG of -0.08 kcal/mol (Table 1, Supplementary Figure 2D). A negative change in ΔΔG suggests that both mutations could result in destabilization of corresponding proteins, signifying distinct downstream consequences of ADAR editing between the two patient groups. Furthermore, overrepresentation analysis (ORA) of genes associated with unique edits revealed significant enrichment (padj <0.05, BH q-Value < 0.05) of distinct biological processes and pathways. Notable, unique edits in non-critical patients were spread across a broader range of biological processes and pathways compared to critical patients. These results suggest that these unique editing sites might have distinct functional roles in influencing COVID-19 disease severity (Fig. 2J, K and Supplementary Table 2E, F).

**Table 1:** Shows the position and potential consequences of non-synonymous substitution within DHRSX gene (unique to critical patients) and HLA-DRB5 (unique to non-critical patients), on corresponding protein stability as predicted by DDMut (Zhou et al., 2023). (H= Histidine; R= Arginine; S= Serine; G= Glycine; ΔΔG = Change in Gibbs Free Energy).

| Disease Severity | Gene name | Exon/position | Wildtype residue | Mutant residue | Predicted stability change ($\Delta\Delta G^{\text{stability wt mt}}$) |
| --- | --- | --- | --- | --- | --- |
| Critical | DHRSX<br>(dehydrogenase/reductase X-linked) | Exon 7; 292 | H | R | -1.27 kcal/mol |
| Non-critical | HLA-DRB5 (major histocompatibility complex, class II, DR beta 5) | Exon 3; 164 | S | G | -0.08 kcal/mol |

### Filtering high confidence ADAR editing sites and identification of differentially edited sites

From the total ADAR editing sites, we further identified a list of high confidence ADAR editing sites using a multistep filtering approach. First, we retained only those A-to-G and T-to-C substitutions that were present in the variant calling outputs of both GATK and JACUSA2. To this list, additional filters were applied on the total number of aligned reads and editing levels per site editing (refer to Methods for details). Additionally, only those sites that were present in at least 20% samples were selected. Following this filtering, a total of 13,531 high confidence ADAR editing sites were identified within 2976 genes (Fig. 3A, Supplementary Table 3A). Of these high confidence ADAR editing sites, 12,828 sites were previously identified, confirmed editing sites present in REDIportal database (Supplementary Table 3B), while the remaining 703 were not present in REDIportal (Supplementary Table 3C), and will be referred to as novel editing sites (Fig. 3B).Moreover, to ensure the robustness of the identified high confidence sites, we examined these sites for features commonly associated with ADAR edited sites. Previous studies have shown that ADAR editing sites are mainly located in the non-coding, particularly within the intronic and 3’UTR regions of the transcripts (Bahn et al., 2012; Hundley & Bass, 2010) and show sequences preferences (motifs) surrounding the edited base (Piontkivska et al., 2021). Consistent with this, both known and novel high confidence sites were enriched in the 3’UTR and intronic regions (Fig. 3C, Supplementary Table 3A). Furthermore, for both known (sites present in REDI portal) and novel high confidence sites, edited A’s on the positive strand had a 5’ neighboring T, C, or A and a 3’ neighboring G (Fig. 3D, E). In contrast, edited T (=U, A on the complementary strand) had a 3’ neighboring A, G, or T (U), or a 5’ neighboring C (Fig. 3F, G).

**Figure 3.**
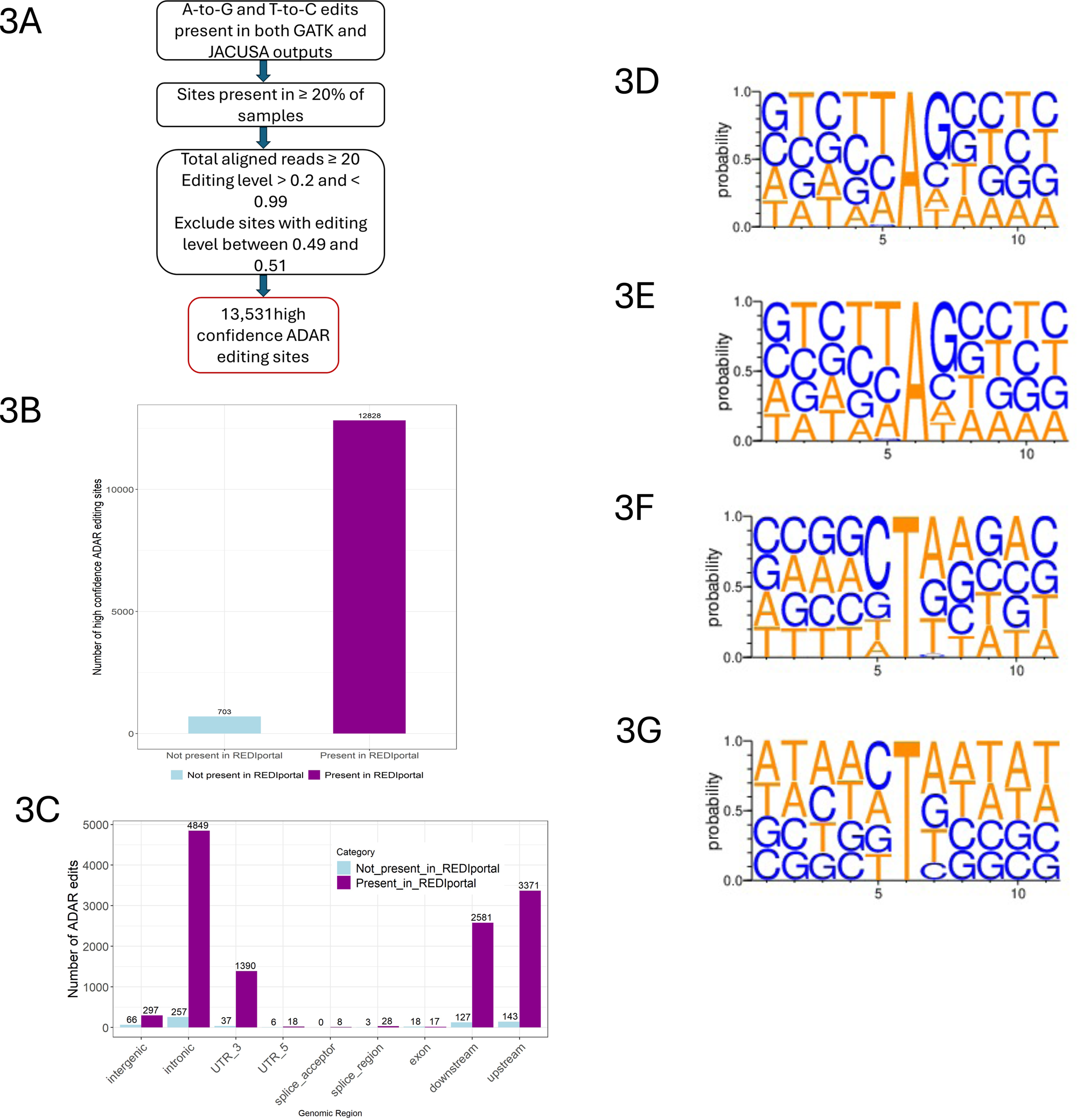
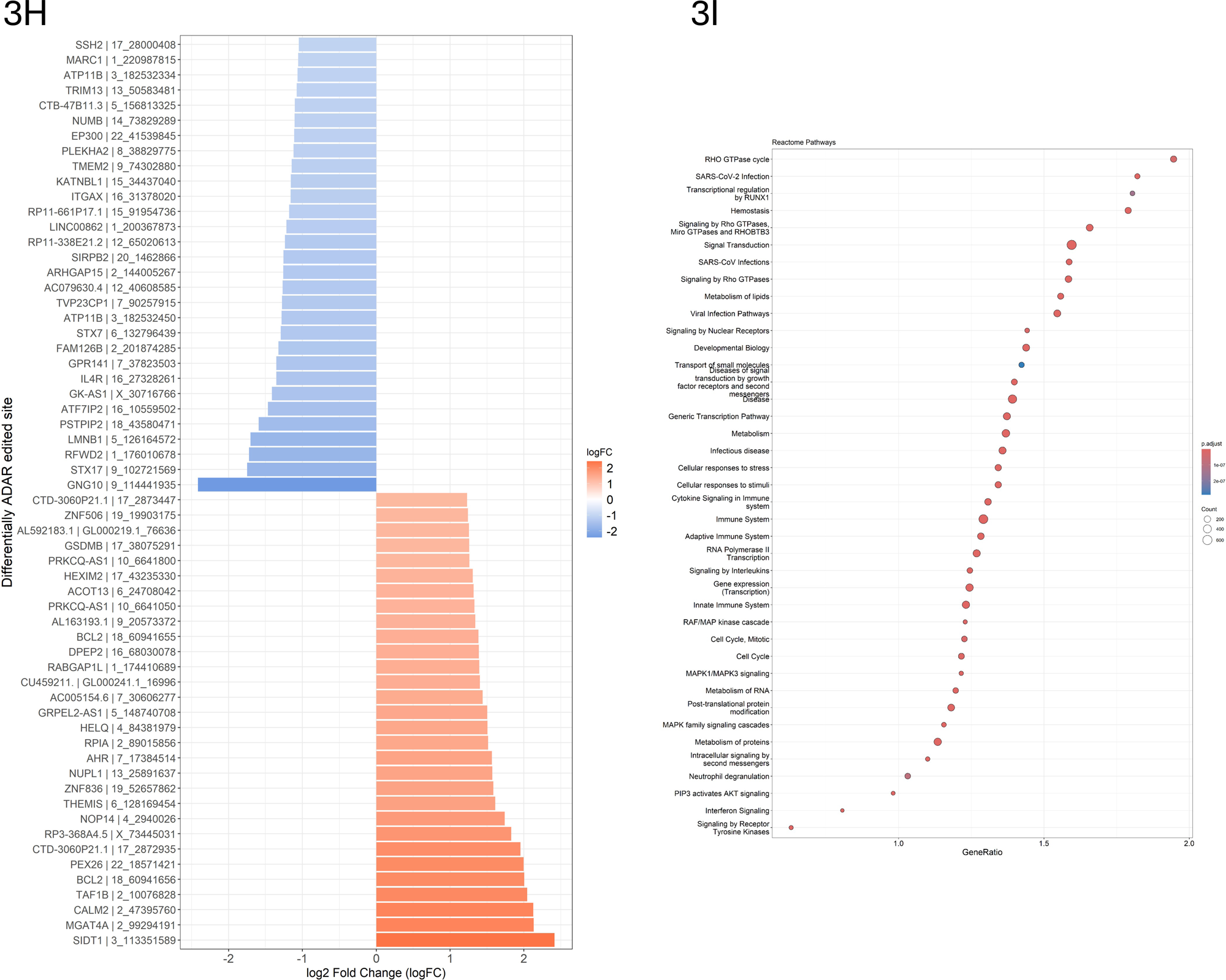
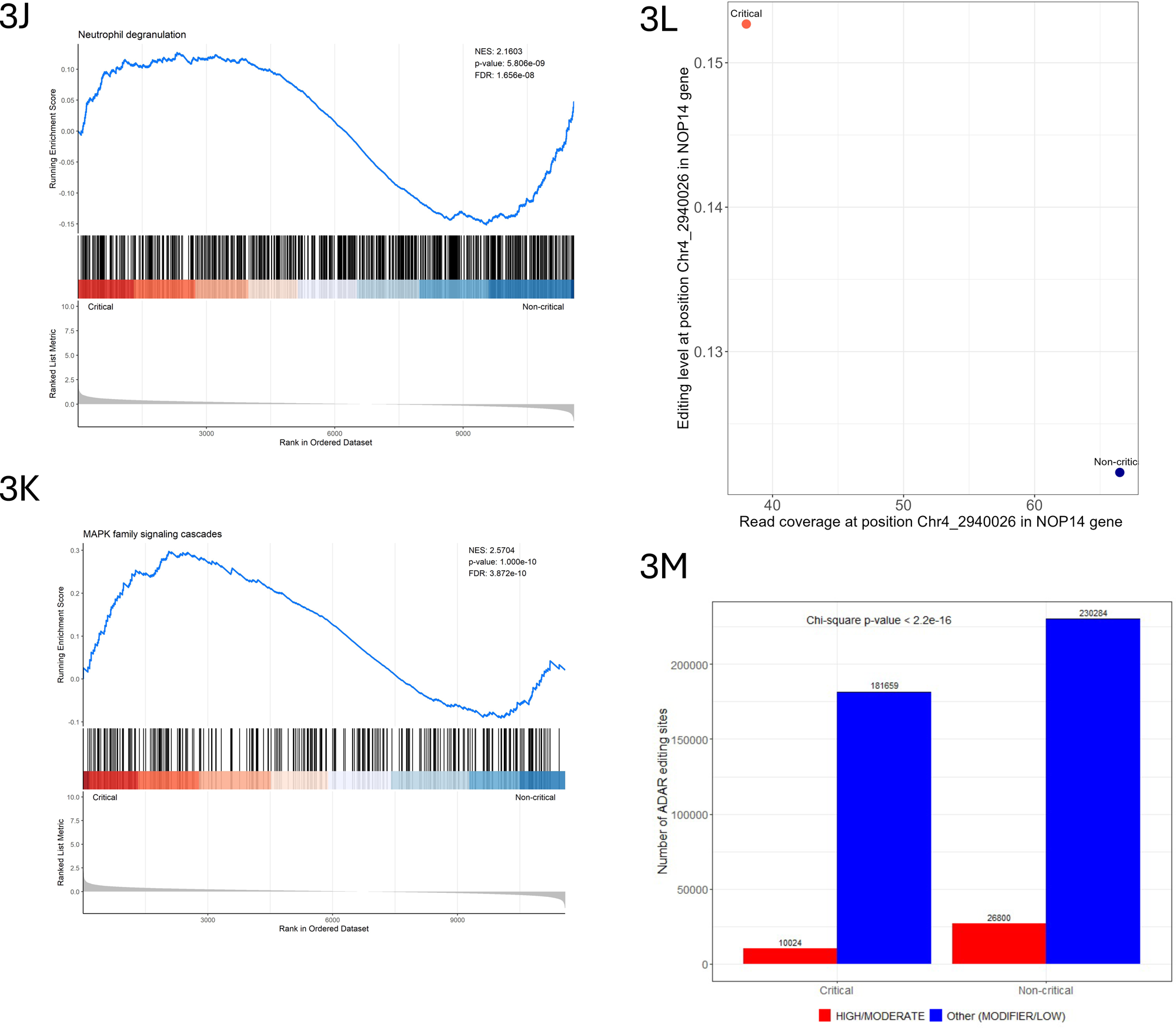
High confidence ADAR editing sites and differential editing. (A) Flow chart of methods/tools and filters used to identify high confidence sites. (B) Bar plot showing the number of high confidence ADAR editing sites that are present in REDIportal, representing previously identified ADAR editing sites, and novel sites, not present in REDI portal. (C) Genic distribution of both known and novel high confidence edits. Sequence logo for edited A’s on the positive strand for both known (D) and novel (E) editing sites. Sequence logo for edited T (=U, A on the complementary strand) for both known (F) and novel (G) editing sites. (H) Bar plot of top 30 sites with significantly increased (Log2 Fold Change (Log2FC) > 0.58 and FDR < 0.05) and top 30 sites with significantly decreased (Log2FC < -0.58 and FDR < 0.05) editing in critical compared to non-critical patients. (I) GSEA results showing Reactome pathways enriched within genes containing differentially edited sites. GSEA plots show positive enrichment of neutrophil degranulation (J) and MAPK family signaling cascade (K) both positively enriched in critical patients. (L) Shows relationship between read coverage and editing levels (values represented as median) at position Chr4:2940026 within NOP14 gene (M) Differences in the number of high and moderate impact editing events between critical and non-critical patients.

We further aimed to identify differentially edited sites within the list of high confidence ADAR editing sites i.e., sites with significant difference in RNA editing levels between critical and non-critical patients using the method previously described in Riemondly et al., 2018 (Riemondy et al., 2018). Briefly, a generalized linear model (GLM) was applied to each high confidence site, using edgeR, to find editing sites where the edited base (G/C) counts show significant differences between critical and non-critical patients, while controlling for unedited bases (A/T) within each sample. Using this approach, we identified a total of 140 differentially edited sites in 126 genes (edgeR Log2FC > |0.58| and FDR < 0.05) within critical compared to non-critical COVID-19 patients (Supplementary Table 3D). Figure 3H shows the top 30 sites with significantly increased (Log2FC > 0.58 and FDR < 0.05) and top 30 sites with significantly decreased (Log2FC < -0.58 and FDR < 0.05) editing in critical compared to non-critical patients. As expected, a majority of these sites were within the Intronic (35%), 3’UTR (12%), and downstream regions (19%) of the transcript (Supplementary Figure 3A). Importantly, differentially edited sites within NOP14 gene at position Chr4:2940026 (with the higher levels of editing in critical samples - edgeR Log2FC 1.7398 and FDR < 0.01) were identified as a moderate impact, missense variant (I779V) by SnpEff and ANNOVAR (Supplementary Table 3D). We further examined whether this difference in editing within NOP14 is associated with differences in expression of its gene. Interestingly, in critical samples despite low coverage at the edited site, editing levels were higher in critical samples, while non-critical samples had higher coverage at the edited position with low editing levels (Fig. 3L). Our results show that missense editing within NOP14 gene is not simply a consequence of increased expression and that editing adds another layer of regulation in COVID-19 disease severity.

Since editing within 3’UTRs could alter miRNA targeting (Tomaselli et al., 2013), we examined whether the differentially edited sites in 3’UTRs are within known miRNA target regions using data from TargetScan (McGeary et al., 2019). The results showed that except for the site Chr4:39551031 within 3’UTR of SMIM14 gene, all other sites were found to be in the known miRNA targeting regions. Specifically, 15 differentially edited sites within 12 genes were predicted to be targeted by 57 miRNAs (Table 2), suggesting that differences in editing between critical and non-critical patients might also affect miRNA-mediated regulation in the two patient groups.

**Table 2.**
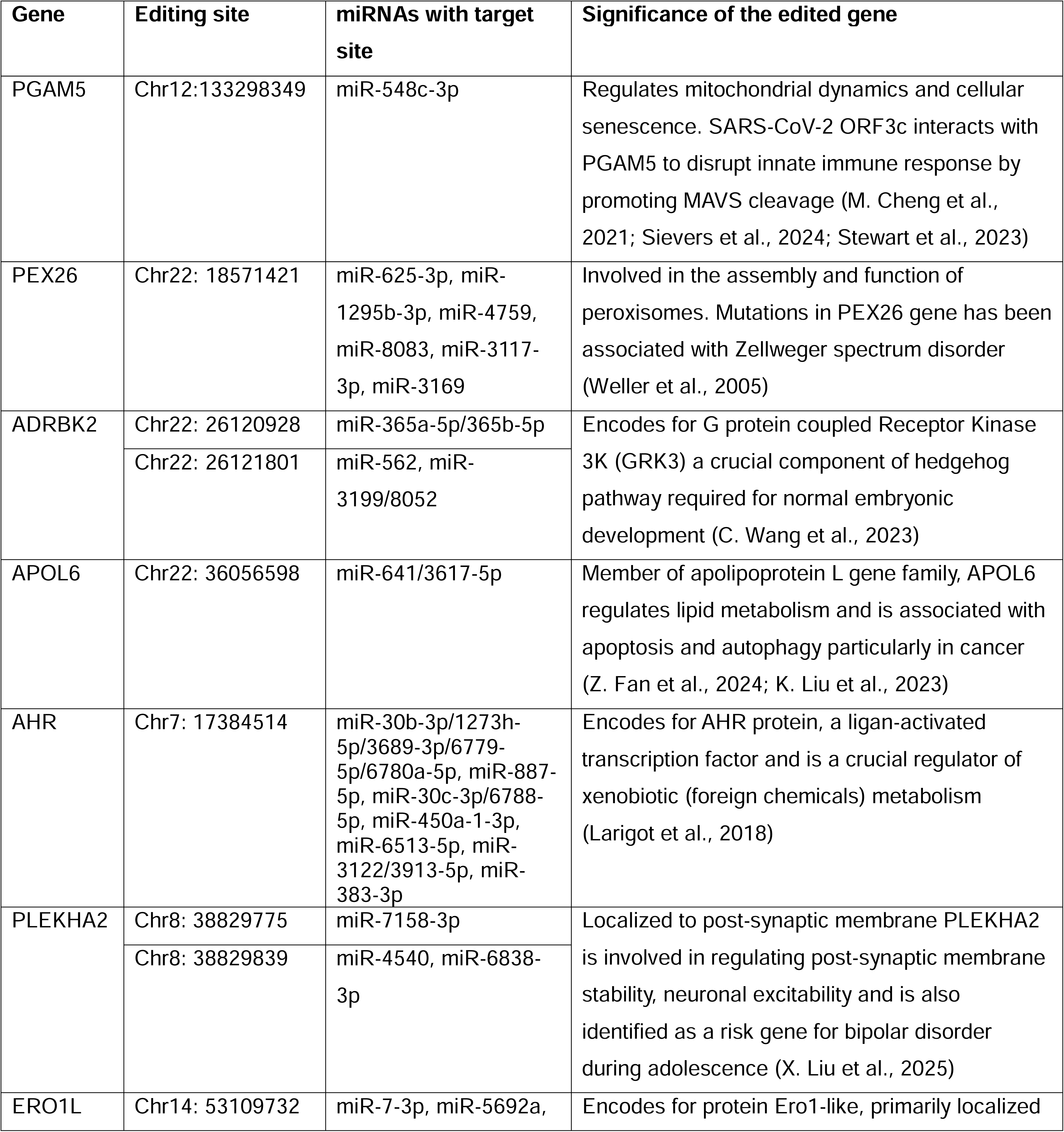
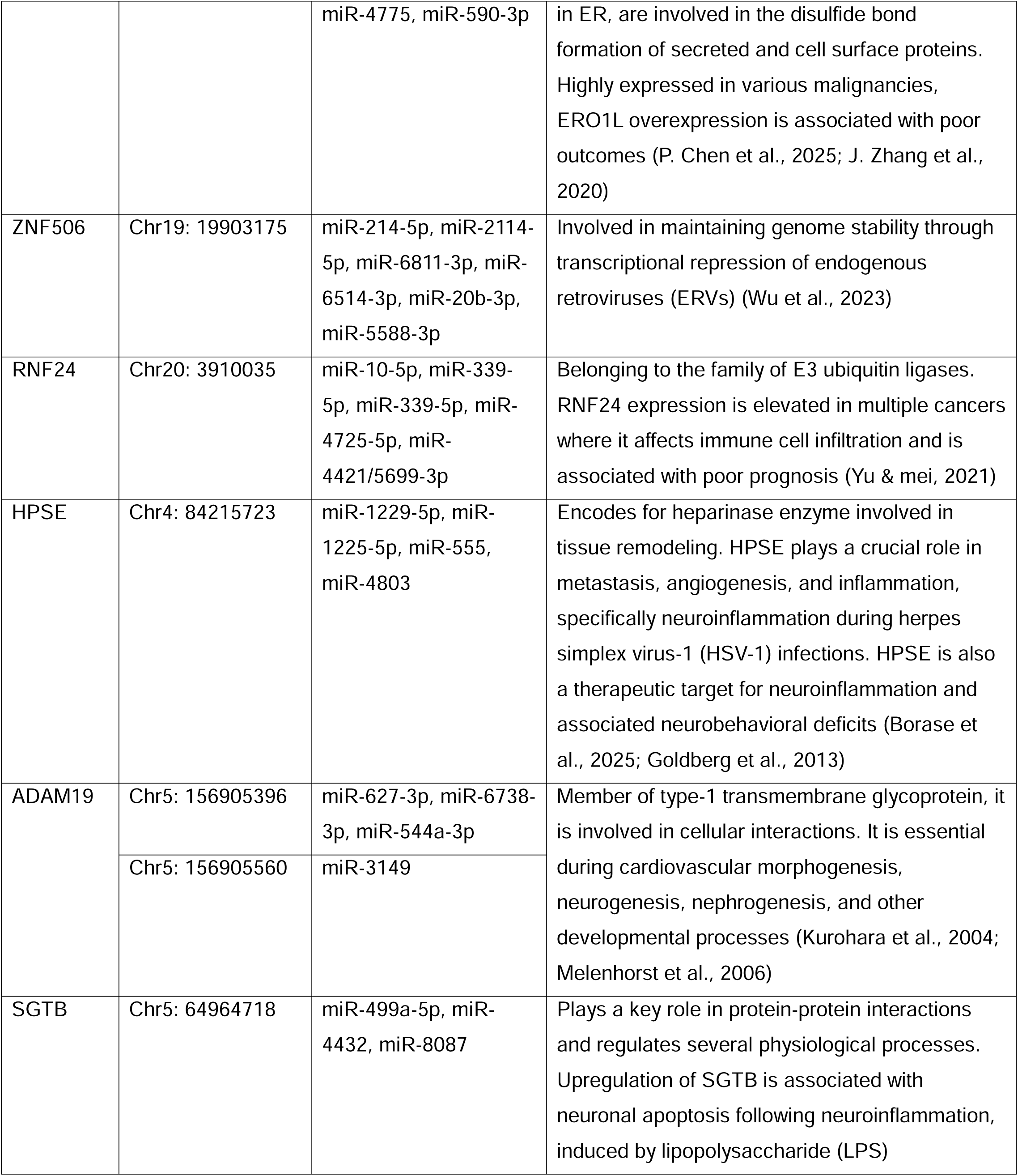
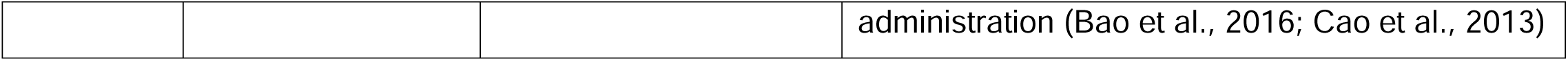
List of differentially edited genes with corresponding ADAR editing sites located within 3’UTR, the microRNAs predicted to target these regions, and significance of the edited genes.

GSEA analysis on a unique list of genes and coordinates (since same genes could have multiple editing site) ranked by Log2FC, identified significant enrichment of multiple key pathways. Notable pathways included, immune system pathways such as “neutrophil degranulation”, “interferon signaling”, “cytokine signaling”, “signaling by interleukins”, and “activation of NF-κB in B cells”. Additionally, infectious disease pathways including “SARS-CoV-2 infection”, signal transduction pathways such as “MAPK family signaling cascade”, “signaling by Rho GTPases, Miro GTPases, and RHOBTB3”, and “insulin receptor signaling cascade” were enriched. Differentially edited genes were also enriched in pathways related to cell cycle, cellular response to stress, and RNA and protein metabolism (Fig. 3I). GSEA plot shows distribution of differentially edited genes within the ranked list for “neutrophil degranulation” pathway (Fig. 3J), and “MAPK family signaling cascade” (Fig. 3K) that plays a key role in inflammation and show positive enrichment in critical COVID-19 patients (Cusato et al., 2023; E. McKenna et al., 2022; Reusch et al., 2021). Interestingly, differentially edited genes were also enriched in pathways associated with cardiac conduction, and neuronal system pathways such as “neurotransmitter receptors and postsynaptic signal transmission” (Supplementary Table 3E). This suggests that ADAR editing may also contribute to cardiac and neurological manifestations observed in patients with critical COVID-19 as reported in previous studies and also identified in some patients within the present dataset (Carapito et al., 2022, Table 1).

### High and moderate impact editing events differ between critical and non-critical COVID-19 patients

SnpEff tool (Cingolani et al., 2012) predicts the functional effect and provides assessment of the putative impact of a variant, classifying it as high, moderate, modifier, and low. Here high/moderate impact editing events represent those editing events that can result in amino acid substitutions likely affecting the structure and function of the encoded protein. On the other hand, modifier/low impact editing events represent those within non-coding regions with insignificant effect on downstream protein (https://pcingola.github.io/SnpEff/snpeff/inputoutput/#eff-field-vcf-output-files). To evaluate differences in the potential functional impact of ADAR mediated RNA editing, we annotated the putative editing sites using SnpEff. Only those A-to-G and T-to-C sites with read depth >= 20 and shared across 20% of samples across a disease severity were selected. We found statistically significant differences (χ^2^ test, p value < 2.2e-16) in the number of high and moderate impact ADAR editing sites, with a significantly lower number in patients with critical COVID-19 (Fig. 3M).

### Random forest classification identifies differentially edited gene signatures associated with COVID-19 disease severity

Next, to identify whether differentially edited sites can differentiate between critical and non-critical patients, we applied a random forest (RF) classifier. The dataset was randomly split to 70% training and 30% test dataset, and the RF model was trained on the training dataset using parameters described in the methods. Model performance was evaluated using receiver operating characteristics (ROC) curve analysis. The classifier showed high performance with an area under the curve (AUC) of 1 for the training dataset (Fig. 4A) and 0.974 on the test dataset (Fig. 4B), indicating that the model is effective in distinguishing between critical and non-critical patients and may be generalized to other datasets. To further identify key editing sites contributing to this classification, we calculated Mean Decrease in Gini Index, a metric commonly used to access the importance of features in RF. The top 30 significant sites ordered according to decreasing Gini Index are shown in figure 4C. Additionally, to assess the relevance of each site in distinguishing disease severity, per site AUC was computed. This resulted in 11 editing sites with AUC values > 0.75 (Fig. 4D/Supplementary Table 4), suggesting fair power (Çorbacıoğlu & Aksel, 2023) of these sites to discriminate between critical and non-critical patients.

**Figure 4.**
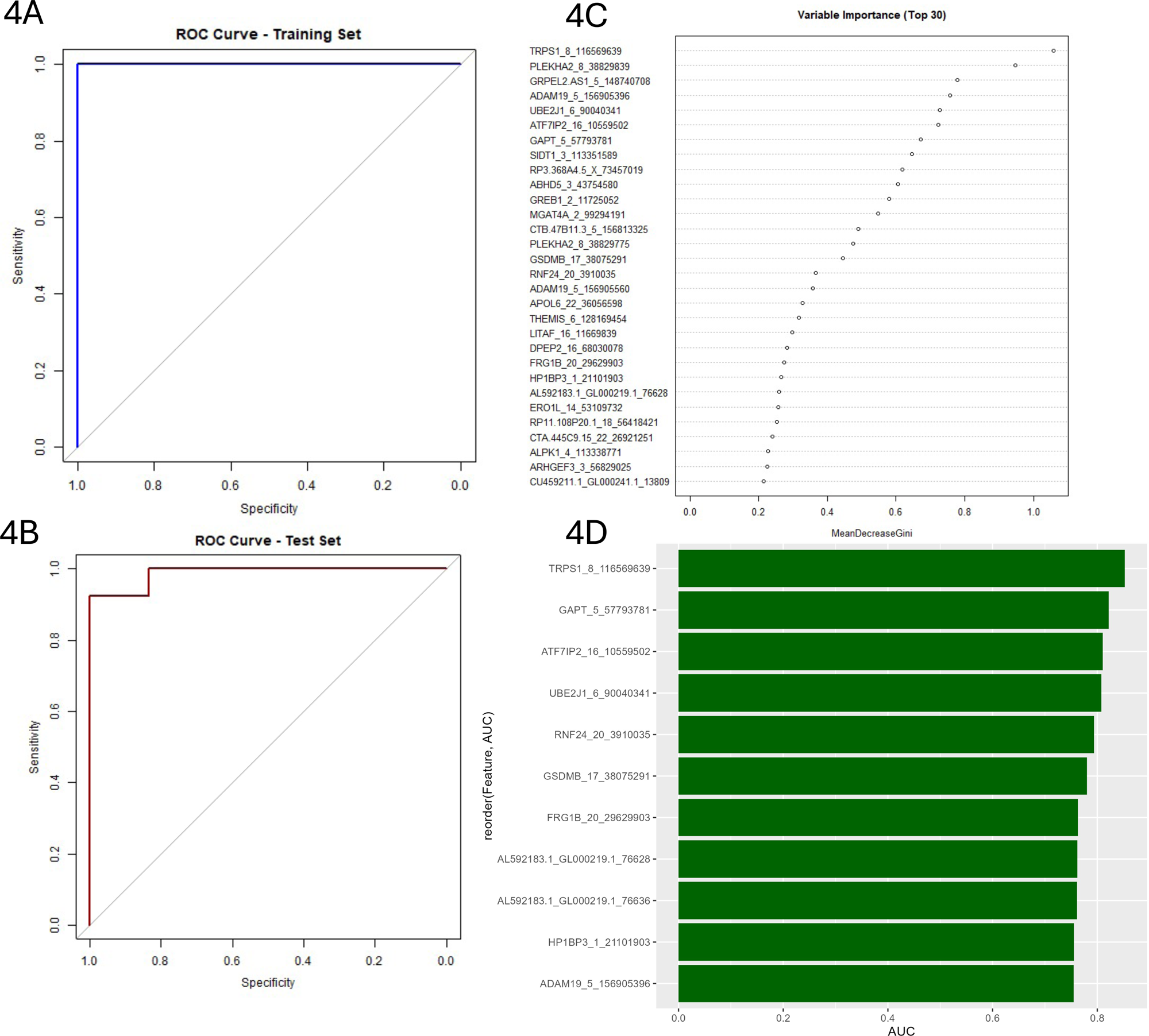
Random forest model identifies RNA editing sites predictive of disease severity. ROC curves showing the performance of random forest classifier in distinguishing critical from non-critical patients in training (A) and test (B) dataset, based on editing levels of differentially edited sites. (C) Variable importance plot showing the top 30 RNA editing sites ranked by Mean decrease in Gini index. (D) Bar plot of top RNA editing sites with AUC > 0.75.

## Discussion

COVID-19 presents with a wide spectrum of clinical outcomes, resulting from complex interplay of various factors, including age, comorbidity status, host genetics, and immune responses (Adab et al., 2022; Biswas et al., 2020; Qian Zhang et al., 2020). Impaired IFN signaling and dysregulation of ISGs is a hallmark of critical disease (Hadjadj et al., 2020; Smith et al., 2022). Among ISGs, ADAR1 encodes for post transcriptional RNA editing enzyme that regulates transcriptome diversity and immune responses, and has been mechanistically linked to symptoms observed both during and post-viral infections (Jin, Liang, Huang, et al., 2024; Piontkivska et al., 2019; Tsivion-Visbord et al., 2020). Previous studies have shown that SARS-CoV-2 infection induces changes in host ADAR expression and editing patterns, contributing to COVID-19 associated physiological changes (Crooke et al., 2021a; Merdler-Rabinowicz et al., 2023; Nair & Piontkivska, 2025). However, to the best of our knowledge, no studies have identified whether the expression and activity of ADARs differ with varying severities of COVID-19.

In this study we address this gap by analyzing a deeply sequenced whole blood RNA sequencing dataset from a comparatively homogeneous cohort of individuals who developed critical COVID-19 and compared them to those who developed non-critical COVID-19, examining differences in ADAR editing patterns in response to SARS-CoV-2 infection. Our analysis revealed significantly high expression of ADAR1 and its isoform ADARp110 in patients with critical COVID-19. We further demonstrate that ADAR activity varies according to disease severity with marked differences observed in 1) the total number of ADAR mediated edits (driven by expression of ADAR1 and ADAR2 in critical but not non-critical patients), 2) editing within repetitive Alu elements, 3) genomic distribution of edited sites, 4) the specific site being edited, including non-synonymous edits predicted to affect protein stability, 5) site-specific editing levels (that within 3’UTR may influence miRNA mediated regulation), and 6) proportion of edits with varying functional consequences. Functional enrichment analysis of uniquely and differentially edited genes in critical patients exhibited significant overrepresentation of inflammatory pathways, including neutrophil degranulation, which was also enriched among differentially expressed genes. Finally, we identified a set of differentially edited sites that could serve as molecular markers of COVID-19 disease severity. Together, our study demonstrates varying expression and editing patterns of ADARs between critical and non-critical patients and proposes a functional role for RNA editing in COVID-19 disease severity.

Extensive differences in the gene expression, including in genes involved in innate immune and inflammatory pathways, particularly neutrophil degranulation, is a key feature distinguishing patients with critical from those with non-critical COVID-19 (Aschenbrenner et al., 2021; Gómez-Carballa et al., 2022), as was observed in the analysis of our dataset as well. While expression of ADAR1, specifically the ADAR1 isoform ADARp110 was significantly higher in patients with critical COVID-19, expression of IFN inducible isoform, ADARp150 did not differ significantly between the two patient groups. Considering that we are comparing virus infected groups (i.e., disease samples) that will result in the production of IFN and ISGs, insignificant differences in ADARp150 levels are understandable. However, it should be noted that the levels of IFNs and ISGs, including ADARp150 are known to be highly dynamic throughout the course of viral infection, and factors such as the timing of sample collection can significantly influence the measurements of expression levels (Chiale et al., 2022). Previous evidence from animal models have shown that the IFN response often peaks a few days after infection and may rapidly decline thereafter (Chiale et al., 2022; Greene & Zuniga, 2021). Therefore, it is possible that ADARp150 levels were different at some point between the two groups of patients; however, further studies that could measure the levels across different time points of SARS-CoV-2 infection are needed to confirm this. Interestingly, ADARp110, although constitutively expressed, is seen to be elevated during viral infections (Nachmani et al., 2014; Tariq & Piontkivska, 2024; K. Zhang et al., 2022), including SARS-CoV-2 infections (Nair & Piontkivska, 2025; X. Peng et al., 2022). This could be due to leaky ribosomal scanning downstream of IFN induced ADARp150 start codon, which results in coexpression of both ADARp110 and ADARp150 (Sun et al., 2021b). It has also been proposed that persistence of SARS-CoV-2 viral transcripts could also drive sustained transcription and translation of ADARp110, resulting in elevated ADARp110 levels (B. Fan et al., 2024). In line with this, an original study by Carapito et al. 2022 (Carapito et al., 2022), reported the presence of viral gene transcripts within samples from patients with critical COVID-19, but not in samples from non-critical COVID-19, which could also be a potential explanation to the higher expression of ADARp110 observed in patients with critical COVID-19 in our analysis.

When comparing differences in global ADAR editing, measured as differences in the total number of ADAR edits, between the two patient groups (comparison between patient groups) we found a lower number of edits within critical patients, despite significantly high expression of ADAR1, compared to non-critical patients. This is consistent with previous studies that have observed that the expression of ADAR1 does not always translate to global higher editing (Deffit & Hundley, 2016; Schaffer et al., 2020; Silberberg et al., 2012). Nonetheless, when we applied a linear regression model to identify relationship between expressions of all three ADARs and the total number of ADAR edits within each patient group (within group correlations), we found significant positive correlation between total number of ADAR edits and the expression levels of ADAR1 and ADAR2 within patients with critical COVID-19. In contrast, no significant correlations were observed between expression of any of the three ADARs and total number of edits for patients with non-critical disease. These results suggest that the activity of ADARs differ between the two patient groups, and that ADAR expression might have a stronger influence on total editing in critical patients, while in non-critical patients factors other than editing might be contributing more to the global editing landscape. In fact, several other factors have been known to influence ADAR activity, including the formation of heterodimers between different ADARs (Cenci et al., 2008), availability of cofactors such as inositol hexaphosphate (IP6) (Macbeth et al., 2005), post transcriptional regulation by miRNAs, post translational SUMOylation (which can reduce ADAR1 editing activity) (Desterro et al., 2005), and availability and accessibility of substrates (Barbon et al., 2007; Bass, 2002). It is also worth noting that ADAR3 is predominantly expressed in neuronal tissues, and is known to have inhibitory effect on the activity of the other two ADARs (Mladenova et al., 2018; Y. Wang et al., 2019), though in our whole blood dataset ADAR3 expression was negligible and showed no correlation with the total number of edits in both groups of patients. It remains possible that ADAR3 may have an impact on editing in neuronal tissues, and further studies using neuronal samples are needed to explore its role in this context. Furthermore, while the total number of ADAR editing events (ie events within both coding and non-coding regions) were lower in critical patients, AEI, a measure of ADAR editing activity within Alu repetitive elements in the genome, was higher in patients with critical COVID-19. Given that ADAR1 is the primary enzyme responsible for editing within Alu elements (Tan et al., 2017; Zhan et al., 2023), and that the expression of ADAR1 was elevated in critical patients, this may explain the observed increase in AEI in critical patients despite lower total number of edits.

In addition to global differences in editing in terms of total number of edits, differences were also seen in ADAR targets, as seen by the presence of genes and pathways that were uniquely edited in either critical or non-critical patients, establishing COVID-19 severity-specific ADAR activity. For instance, non-synonymous editing within exon 7 of the Dehydrogenase/Reductase X-Linked gene (DHRSX), that was predicted to result in protein destabilization, was preferentially edited in patients with critical COVID-19. DHRSX encodes for an enzyme involved in dolichol synthesis, a lipid essential for N-glycosylation (Kentache et al., 2024; Wilson et al., 2024). SARS-CoV-2 spike (S) protein and the host receptor protein angiotensin converting enzyme 2 (ACE2) undergo N-glycosylation facilitating the binding between the two. Considering the importance of N-glycosylation in SARS-CoV-2 infection and the essential requirement of DHRSX gene in N-glycosylation (Das et al., 2024; Zhao et al., 2020), it can be speculated that missense editing in this gene can impact viral entry or replication. While this could be a potential mechanistic link between DHRSX editing and disease severity, further studies are required to confirm this speculation, if any. Similarly, a unique edit was found at position 164 on exon 3 in the gene Human leukocyte antigen class II heterodimer β5 (HLA-DRB5). HLA-DRB5 is a Class II major histocompatibility complex (MHC) molecule involved in antigen presentation to CD4+ T cells and is a key player in adaptive immune responses. Although the direct effect is unknown, when the role of HLA in antigen presentation is considered, it is plausible that non-synonymous editing within this gene may affect viral recognition by the host (Ye et al., 2024). Noteworthy, the extent of protein destabilization resulting from these non-synonymous edits were relatively different, with much larger change in a ΔΔG resulting from non-synonymous editing within DHRDX gene in critical patients, indicating that editing within DHRSX may have a greater impact on protein stability.

The remaining unique edits were predominantly distributed within the intronic and 3’UTR of transcripts, although the numbers varied between critical and non-critical COVID-19 patients. Such editing within non-coding regions may impact multiple processes regulating gene expression and ultimately influencing protein function. For instance, editing within introns can result in conversion of 5’ splice donor sites and/or splice 3’ splice acceptor site, potentially altering splicing patterns and leading to the production of alternative protein isoforms (Bass, 2002; Tang et al., 2020; D. Zhang et al., 2024). Splice site editing may also alter mRNA stability, as seen in increased stabilization of FAK transcript, promoting tumor progression in lung adenocarcinoma (Amin et al., 2017). Additionally intronic editing can alter biogenesis of circular RNAs (circRNAs)(Ivanov et al., 2015; Shen et al., 2022). Similarly, editing within 3’UTR has been shown to impact miRNA targeting, either by creating or destroying miRNA targets (Kawahara et al., 2007; Tomaselli et al., 2015). These findings suggest that severity specific differences in ADAR editing, including in both coding and non-coding regions, could contribute to differences in protein stability, as well as in regulatory processes (such as alternative splicing, mRNA stability, and miRNA targeting) between patients with critical compared to non-critical COVID-19.

Considering the dynamic and heterogenous nature of ADAR editing between individuals and disease conditions (Savva et al., 2012b), we identified a set of high confidence ADAR editing events, with signatures consistent with that of ADAR edited sites. While the majority of these were previously reported sites, a subset of these sites were novel, not previously reported sites, and may represent editing events specifically induced by SARS-CoV-2 infection irrespective of disease severity. Notably, we found that many such high confidence sites were differentially edited between patients with critical and non-critical disease, indicating that the site-specific editing rates also varies with COVID-19 disease severity. For instance, we identified differential editing within multiple genes related to SARS-CoV-2 infection and disease severity. One such differentially edited site was within the 3’UTR of APOL6 (Apolipoprotein L6) gene that has been previously implicated in immune responses to COVID-19 vaccinations and was proposed to contribute to antiviral defense and immune regulation (Jin, Liang, Pan, et al., 2024). Our identification of differentially edited sites within APOL6 gene reinforces its significance in SARS-CoV-2 associated immune responses. We also detected a differentially edited site within DDX6 (DDX DEAD box RNA helicase 6), an RNA helicase involved in RNA metabolism and p-body formation. DDX6 has been shown to be essential for SARS-CoV-2 replication. The viral nucleocapsid (N) protein is known to interact with DDX6 and hijacks it to promote viral replication. This suggests that differential editing within DDX6 may influence its function or regulation and impact SARS-CoV-2 replication thereby impacting disease severity (Ariumi, 2022). Furthermore, several intronic edits were found within the BCL2 (B-Cell Leukemia/Lymphoma 2) gene, a key anti-apoptotic gene from the intrinsic (mitochondrial) apoptotic pathway. Notably, low levels of BCL2 have been reported in patients with critical disease (specifically in non-surviving patients), which is associated with higher cellular damage due to apoptosis. Although the functional consequences of intronic editing remain unclear, it is possible that such edits may influence BCL2 expression or splicing (Lorente et al., 2021). We also identified a differentially edited site within NOP14 gene that results in a non-synonymous edit. Importantly, this editing event was independent of the expression of the host gene, suggesting that this editing is not simply a consequence of increased expression. Moreover, Peng et al. (L. Peng et al., 2016) had previously identified the exact site to be differentially edited in patients with lung adenocarcinoma, further underscoring the importance of ADAR editing at this site in diseases. Differential editing was also observed in genes previously identified to undergo editing across various diseases. For instance, editing within PGAM5 has been associated with heart diseases (J. Chen et al., 2021), ITGAX has been shown to be differentially edited in children with *Mycoplasma pneumoniae* pneumonia (Jin et al., 2025), NLRC5 has been reported to be edited in the autoimmune condition, primary Sjögren’s syndrome (X. Wang et al., 2023). The consistent editing within disease related genes indicates that differentially edited sites identified in our analysis may hold clinical significance.

Overall, this study takes advantage of a deeply sequenced and relatively homogenous cohort in terms of age and comorbidity status, factors that can impact severity and editing patterns (Biswas et al., 2020; Nicholas et al., 2010; Szymczak et al., 2022), thereby enabling a comparison of differences in ADAR editing across varying severities of COVID-19. Moreover, these patients were sampled during the first wave of the COVID-19 pandemic in France, indicating the infections were potentially due to an ancestral SARS-CoV-2 strain (Carapito et al., 2022). Furthermore, these samples were obtained prior to widespread use of corticosteroids in COVID-19 treatment. Corticosteroids are anti-inflammatory drugs known to reduce mortality and morbidity in critical patients (Barnes, 2006; Van den Eynde et al., 2021). However, these drugs are known to influence ADAR editing rates (Vlachogiannis et al., 2020), therefore, sampling before the use of corticosteroids reduces the risk of treatment related factors confounding the results. We also would like to point out the relatively high sequencing depth of this dataset. The reliability of detected RNA editing events is known to increase with the coverage, and a depth of 80-100 million reads is typically recommended for Illumina sequencing (Diroma et al., 2019). Our dataset had an average depth of ∼208 million reads per sample, sufficient to detect editing even within lowly expressed genes. Nonetheless, despite these strengths, our study also has some limitations. Firstly, host genetics plays a very significant role in deciding COVID-19 disease severity; however, the presence of disease affecting variants, if any, was not known in these patients. Moreover, two patients with critical COVID-19 were reported to have anti-type I IFN autoantibodies in the parent study (Carapito et al., 2022); thus, it is possible that this may influence ADAR expression and downstream editing. The parent study also stated that the presence of obesity alone in the patients was not considered as an exclusion criteria under comorbidity; however, multiple studies have identified obesity as a major comorbidity affecting COVID-19 severity and can also alter editing patterns (Cui et al., 2021; Nagy et al., 2023). Furthermore, it should be noted that ADAR editing is highly variable between tissues, and thus, blood ADAR editing signatures and presumed downstream effects may not fully represent the physiological changes within other tissues, especially those in neuronal tissues where editing is highly enriched and expression of the catalytically inactive enzyme ADAR3 impacts the activity of other two ADARs. Nonetheless, our findings show that patterns of ADAR editing could indeed serve as prognostic and/or diagnostic markers for COVID-19 disease severity.

## Methods

### Transcriptomic dataset of COVID-19 patients with critical and non-critical COVID-19

In this study, we used deeply sequenced transcriptomic data, with an average depth of ∼208 million reads per sample (Supplementary Table 5), from Carapitio et al. study (Carapito et al., 2022), which is publicly available at NCBI GEO under the accession GSE172114 (BioProject PRJNA722046). Briefly, the dataset includes whole blood RNA sequencing samples from 69 young patients (under the age of 50), without comorbidities, hospitalized for varying severities of COVID-19 (confirmed using qRT-PCR test of nasopharyngeal swabs) at a university hospital network in northeast France from March to April 2020 (first wave of pandemic in France). These 69 patients were categorized into two groups based on severities of COVID-19 presentations. The first group consists of 23 patients with non-critical COVID-19, referred to as those who stayed at non-critical a care ward and may or may not have required low flow supplemental oxygen. The second group consists of 46 patients with critical COVID-19 defined as those hospitalized in the Intensive Care Unit (ICU) due to moderate or severe ARDS (berlin criteria) and/or requiring high flow nasal oxygen and mechanical ventilation due to respiratory failure (Table 3). The group also included patients who succumbed to the disease. Detailed summary of patient characteristics including drug and supportive treatments (if used) is available in Carapito et al. (2022), Table 1.

**Table 3:** Characteristics of BioProject PRJNA722046 from Carapito et al. (2022) used in this study. Values for the number of ADAR editing sites are represented as mean and standard error of mean (SEM) for respective severity group.

| COVID-19 disease severity | Sample Count | Average number of aligned reads | Mean number of putative ADAR editing sites (+/- SEM) per sample for different severities of COVID-19 | Mean A-to-G Alu Editing Index (AEI) (+/- SEM) per sample for different severities of COVID-19 |
| --- | --- | --- | --- | --- |
| Non-critical | 23 | 203,697,837 | 158022.21 ± 10234.06 | 0.847 ± 0.0333 |
| Critical | 46 | 197,249,840 | 139285.2174 ± 9307.70 | 0.893 ± 0.0285 |

### RNA-seq data analysis and identification of RNA editing sites

RNA sequencing dataset of paired end 151 base pairs (bp) long reads from total RNA, depleted for ribosomal and globin RNA, from Carapito et al. (2022 (Carapito et al., 2022), were downloaded from NCBI SRA. Quality check, read mapping, and assembly were performed using the computational pipeline Automated Isoform Diversity Detector (AIDD) (Plonski et al., 2020). Briefly, raw FASTQ files were quality-checked using FASTQC (https://www.bioinformatics.babraham.ac.uk/projects/fastqc/). Following trimming, reads were aligned to GRCh37 (human reference genome Ensembl build release 75) annotated with splice sites and genomic single nucleotide polymorphisms using HISAT2 (D. Kim et al., 2015). On average, approximately 200 million reads were aligned to the genome (Supplementary Table 5). Stringtie (Pertea et al., 2015) was used to perform transcriptome assembly, the resulting ballgown files with gene and transcript level information were analyzed using a custom bash script to generate gene and transcript level raw count matrices. Both gene and transcript counts were analyzed using DESeq2 (v1.44.0) (Love et al., 2014) to identify differentially expressed genes, including ADARs, between critical and non-critical patients. Genes that met the criteria of an adjusted p-value (padj) < 0.05 and log2 Fold Change (Log2FC) less than > |0.58| were considered differentially expressed. ADAR isoform expression was measured as DESeq2 normalized counts. FDR correction for multiple testing was performed using the Benjamini and Hochberg (BH) method. Two different tools, Genome Analysis Toolkit (GATK) (A. McKenna et al., 2010) haplotype caller and JACUSA2 (Piechotta et al., 2022) were used to identify all potential RNA editing events. GATK analysis followed the best practices settings as defined in Plonski et al., (Plonski et al., 2020) to identify putative ADAR editing sites from the BAM files. Single nucleotide polymorphisms (SNPs) defined in NCBI database of SNPs were filtered out (Sherry et al., 2001). Java framework for accurate SNV assessment (JACUSA2) (Piechotta et al., 2022), filtering out sites with less than 20 reads, was used to identify putative ADAR editing sites and count the number of bases observed at each such site. ADAR editing events were defined as substitutions that include both A-to-G and U (T)-to-C, the latter representing editing events on the complementary strand, referred here to as T-to-C substitutions. Thus, identified editing site matrixes were annotated, and their potential functional consequences were predicted using snpEff variant annotation and effect prediction tool (Cingolani et al., 2012) and ANNOVAR gene symbols using RefGene (K. Wang et al., 2010).

### Calculation of Alu editing index (AEI)

Alu editing index (AEI) represents the weighted average of all the A-to-G mismatches within Alu elements to the total coverage of adenosines within a genomic region. Alu editing indexer (AEI) method described in Roth et al., 2019 (Roth et al., 2019), using default parameters on STAR mapped BAM files as input, was used to determine differences in editing levels within the Alu repetitive elements, between the two patient groups.

### Selection of high confidence ADAR editing sites and differential RNA editing analysis

High confidence ADAR editing sites were identified using multiple filters on GATK (A. McKenna et al., 2010) and JACUSA2 (Piechotta et al., 2022) generated VCF files. First, GATK and JACUSA2 generated VCF files were filtered to include only those A-to-G and T-to-C substitutions that were identified by both the tools. Further, only those substitutions that were present in at least 20% of all samples and with greater than or equal to 20 total aligned reads (read coverage), editing level (defined as the proportion of G reads at a site with an A reference or C reads at a site with T reference) greater than 0.2, less than 0.99 and not between 0.49 and 0.51. The latter filtration was done to include only sites with at least 20% editing level at each edited site and to exclude potential noise, homozygous and heterogenous genomic variants respectively. The remaining sites were further mapped to REDIportal (Picardi et al., 2017), a comprehensive database providing curated collection of ADAR editing events across multiple tissues including whole blood, to identify known ADAR editing sites. Sites that were not present in REDIportal were classified as novel sites. Next, to identify differentially edited sites between critical and non-critical patients we applied a statistical model using the edgeR package (Version4.2.2) (Robinson et al., 2010), as described previously in Reimondy et al., 2018 (Riemondy et al., 2018). Briefly, for each high confidence ADAR editing site, we extracted the reference and alternative base counts from each sample, generating a count matrix consisting of separate counts for each base per sample. A generalized linear model (GLM) was then constructed, ∼0 + sample_ID + condition:allele, where “sample_ID” accounts for individual specific effects, “condition” represents disease severity, and “allele” indicates reference or alternative base. Dispersion estimates were calculated and the GLM was fit to the above count data. Differential editing analysis was performed using a likelihood ratio test (glmLRT) to identify significant differences in edited base count across conditions, while controlling for within sample differences in reference base count. Differentially edited sites were defined as those with Log2FC > |0.58| and FDR <0.05.

### Enrichment analysis

To identify functionally relevant pathways and biological processes within our gene list, we performed both Over Representation Analysis (ORA) and Gene Set Enrichment Analysis (GSEA) using the R package - “cluster profiler” (Version 4.12.6). ORA was performed on the list of genes unique to disease severity to identify Gene Ontology (GO) terms and Reactome pathways enriched within the unranked list of genes. GSEA was performed on ranked gene lists generated from both, DESeq2 analysis (to identify differentially expressed genes), and edgeR analysis (to identify differentially edited ADAR editing sites). In both analyses, for GSEA, genes were ranked based on Log2FC values, and 1000 permutations were used to access statistical significance. Gene annotations were provided using the org.Hs.eg.db human gene annotation database. Pathways with an adjusted p-value < 0.05 were considered significantly enriched and ordered according to enrichment score.

### Motif Analysis

Both the novel and known high confidence ADAR editing sites (A-to-G and T-to-C analyzed separately) were converted to BED format. BED files were then used with BEDtools (Quinlan & Hall, 2010) to extract 5 bp upstream and downstream sequence flanking the edited bases from the unmasked human reference genome (Homo_sapiens.GRCh37.dna_sm.primary_assembly.fa). The resulting FASTA sequence was then used to generate sequence logos using Weblogo (version 3.7.12) (Crooks et al., 2004) using a custom bash script.

### *In silico* prediction of effect of missense edits on protein stability

DDMut (Zhou et al., 2023), with AlphaFold2 (J. Cheng et al., 2023; Jumper et al., 2021) predicted protein structure as input, was used to predict changes in protein stability caused by the missense edits. DDMut is a deep learning model that predicts changes to protein stability, quantified as changes in Gibbs free energy (ΔΔG) in kcal/mol, based on atomic interactions in the wild type and mutant amino acid residue. A negative ΔΔG value (< 0) indicates protein destabilization while positive ΔΔG (> 0) indicates protein stabilization.

### Analysis of miRNAs with targets overlapping with differentially edited sites within 3’UTRs

Because editing within 3’UTRs can alter miRNA targeting (Tomaselli et al., 2013), we examined whether the differentially edited sites in 3’UTRs are located within known miRNA target regions using data from TargetScan (McGeary et al., 2019). All pre-identified miRNA target sites were obtained from TargetScan (v8.0) by downloading All_Target_Locations.hf19.bed file from https://www.targetscan.org/vert_80/vert_80_data_download/All_Target_Locations.hg19.bed.zip. Differentially edited sites within 3’UTRs were compared to the miRNA target sites using custom R script. Genomic coordinates of 3’UTR editing sites and miRNA target regions were matched using R package GenomicRanges (Lawrence et al., 2013).

### Random forest and ROC analysis

To assess whether ADAR editing could distinguish between critical and non-critical patients, we used a random forest (RF) classifier. The ADAR editing matrix containing editing levels for all the identified differentially edited sites was used as features. Samples were randomly partitioned to test (70%) and training (30%) and the random forest was trained on the training data using randomForest (Liaw & Wiener, 2002) package in R, with 1000 trees default parameters. Model performance was evaluated using receiver operating characteristics (ROC) curve and area under the curve (AUC), computed using the pROC package (Robin et al., 2011). Editing sites that were most important for classification were calculated using the Mean Decrease in Gini index, a measure of variable importance in RF (Nembrini et al., 2018). Additionally, site specific predictive power of a site was calculated using AUC values, where each site was used as a predictor for binary outcome. Editing sites with AUC> 0.75 was considered to have fair discriminatory power (Çorbacıoğlu & Aksel, 2023).

### Statistical analysis of expression and editing data

The Wilcoxon rank-sum test was used to compare the expression of ADAR1 isoforms, ADARp110 and ADARp150, as DESeq2 normalized values between critical and non-critical patients. Pearson’s Chi-squared test with yates’ continuity correction was used to compare the number of High/Moderate or other editing events between the two patient groups. To examine whether ADAR gene expression is associated with the total number of ADAR editing events within each patient group, we performed linear regression analysis using expression values of all three ADARs in transcripts per million (TPMs), as independent variable and total number of ADAR edits within each group as dependent variable. The strength and significance of these associations were assessed using adjusted R^2^ and p-values (<0.05).

Supplementary materials and code are shared at https://github.com/RNAdetective/Editing_in_critical-non-critical-COVID-19.

## List of abbreviations

ADAR: Adenosine Deaminase Acting on RNA
IFN: Interferon
ISG: Interferon stimulated gene
ISRE: Interferon-stimulated response element
UTR: Untranslated region
SARS-CoV-2: Severe acute respiratory syndrome coronavirus 2
COVID-19: Coronavirus disease 2019
ARDS: Acute respiratory distress syndrome
ssRNA: single stranded RNA
dsRNA: double stranded RNA
SINE: Short interspaced repetitive elements
PAMP: Pathogen associated molecular patterns
PCA: Principal component analysis

