## Supplementary material for "SARS-CoV-2 infection induced alterations in ADAR editing patterns differ between patients who developed critical compared to non-critical COVID-19": SupplFiles_ST1-5_SF1-3: Supplementary_figures_1-3.docx

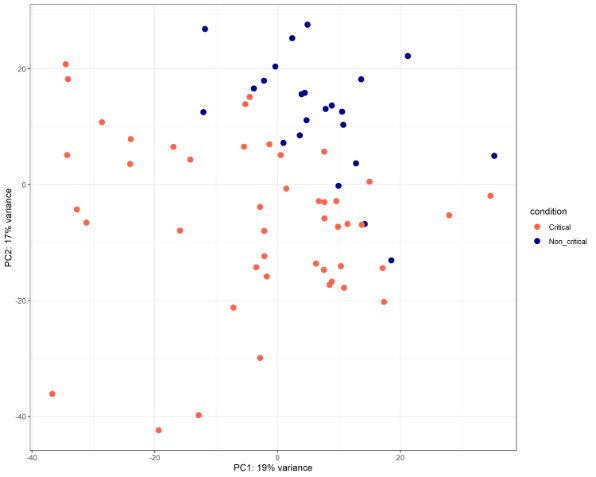

**Supplemental Figure 1:** PCA plot of top 500 differentially expressed genes. PC1 clearly separated critical from non-critical patients.

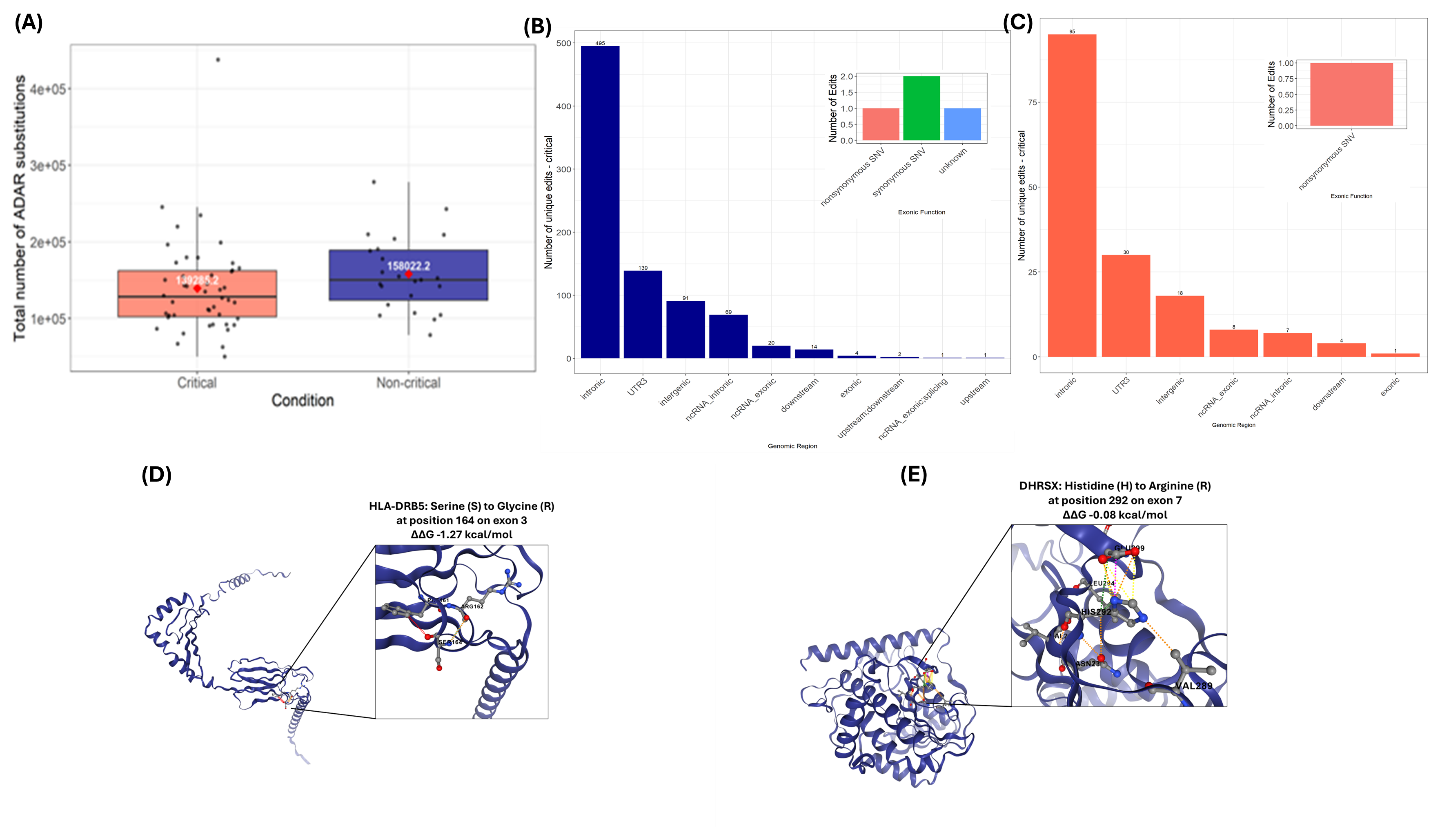

**Supplemental figure 2: Difference in ADAR editing between critical and non-critical patients** (A) Average number of ADAR edits between critical and non-critical patients. (B) Region wise distribution of uniquely edited sites within non-critical patients. inset plots show different types of substitutions within exonic regions. (C) Region wise distribution of uniquely edited sites within critical patients. inset plots show different types of substitutions within exonic regions. (D) Screenshot of interactive 3D viewer showing interatomic interactions of the wildtype residue (S) at position 164 in exon 3 in HLA-DRB5 gene carrying non-synonymous substitution in non-critical patients. (E) Screenshot of interactive 3D viewer showing interatomic interactions of the wildtype residue (H) at position 292 in exon 7 in DHRSX gene carrying non-synonymous substitution in critical patients.

|  | Total |
| --- | --- |
| 3_prime_UTR_variant | 16 |
| downstream_gene_variant | 27 |
| intergenic_region | 4 |
| intragenic_variant | 2 |
| intron_variant | 49 |
| missense_variant | 1 |
| upstream_gene_variant | 41 |

**Supplemental Figure 3:** Genomic distribution of differentially edited sites. Highest number of differentially edited sites mapped to introns followed by 3’UTR, exonic region also contains a missense edit.
